# Nucleus translocation of tRNA synthetase mediates late integrated stress response

**DOI:** 10.1101/2020.06.07.138792

**Authors:** Na Wei, Haissi Cui, Yi Shi, Guangsen Fu, Navin Rauniyar, John R. Yates, Xiang-Lei Yang

## Abstract

Various stress conditions are signaled through phosphorylation of translation initiation factor eIF2α to inhibit global translation while selectively activating transcription factor ATF4 to aid cell survival and recovery. However, this integrated stress response is acute and cannot resolve lasting stress. Here we report that TyrRS, a member of the aminoacyl-tRNA synthetase family capable of responding to diverse stress factors through cytosol-nucleus translocation and activating stress-response genes, also inhibits global translation, however at a later stage than eIF2α/ATF4 and mTOR responses. Excluding TyrRS from the nucleus over-activates protein synthesis and increases apoptosis in cells under prolonged oxidative stress. Nuclear TyrRS transcriptionally represses translation genes by recruiting TRIM28 and/or NuRD complex. We propose TyrRS, possibly along with other family members, can sense a variety of stress signals through intrinsic properties of this enzyme and its strategically located nuclear localization signal and integrate them by nucleus-translocation to effect protective responses against prolonged stress.

## Introduction

Regulation of mRNA translation is a key component of cellular stress response. Two major pathways - the regulation of mammalian target of rapamycin (mTOR) and the phosphorylation of initiation factor eIF2α - are well studied for translation control at the initiation step (Sonenberg and Hinnebusch, 2009; Jackson et al., 2010; Spriggs et al., 2010). Hypoxia and energy depletion disable eIF4E, via the regulation of mTOR, to prevent cap-dependent translation initiation (Kim et al., 2002; Marcotrigiano et al., 1999; Reiling and Sabatini, 2006). In response to a wide variety of stresses, both extrinsic factors such as amino acid deprivation and viral infection or intrinsic conditions such as endoplasmic reticulum (ER) stress caused by accumulation of unfolded proteins, a common adaptive pathway, termed the integrated stress response (ISR), is activated to reduce the availability of translation initiation ternary complex (eIF2-GTP-Met-tRNA^Met^) via the phosphorylation of eIF2α by a family of upstream kinases (Shenton et al., 2006). These two pathways are largely responsible for the acute reduction of protein synthesis in cells under stress conditions.

However, as a prolonged decrease in translation would be harmful, cells also subsequently trigger negative feedback loops that reactivate protein synthesis. Indeed, the ISR-induced phosphorylation of eIF2α inhibits global translation while selectively enable the translation of the activating transcription factor 4 (ATF4) and CCAAT/enhancer-binding proteins homologous protein (CHOP). In addition to promoting the transcription of stress response genes, ATF4 and CHOP also promote the expression of protein phosphatase 1 regulatory subunit 15A (PPP1R15A/GADD34), which dephosphorylates eIF2α to reinstate physiological protein synthesis (Harding et al., 2000; Novoa et al., 2001). Moreover, ATF4 and CHOP interact to directly induce genes encoding protein synthesis (Han et al., 2013). In cells with unresolved stress, the ATF4/CHOP-mediated activation of protein synthesis causes ATP depletion and additional reactive oxygen species (ROS) production, leading to cell death (Han et al., 2013; Marciniak et al., 2004). Thus, the eIF2α-ATF4/CHOP feedback loop prevents over-suppression of protein synthesis under stress but can also trigger cell death when the stress condition is not resolved quickly.

The largest gene family targeted by ATF4 and CHOP related to translation is aminoacyl-tRNA synthetases (aaRSs) (Han et al., 2013). These cytoplasmic enzymes catalyze an ATP-dependent tRNA aminoacylation reaction, which provides building blocks for ribosomal synthesis of growing peptides according to the genetic code. Interestingly, many aaRSs are also found in the nucleus of eukaryotic cells where translation generally does not occur. The initial hypothesis was that aaRSs function here in proofreading newly-synthesized tRNAs (Hopper and Huang, 2015; Lund and Dahlberg, 1998). However, later findings suggest that the nuclear-localized aaRSs are involved in regulating a wide range of biological processes including vascular development, inflammation, energy production, and stress responses, mainly due to their distinctive abilities to interact with the transcriptional machinery and DNA (Frechin et al., 2014; Shi et al., 2014; Wei et al., 2014; Yannay-Cohen et al., 2009).

Interestingly, when the nuclear localization signal (NLS) in human tyrosyl-tRNA synthetase (TyrRS) was identified, we noticed that the NLS is directly involved in tRNA binding at the anticodon (Fu et al., 2012). As a result, the nuclear translocation of the synthetase is negatively regulate by its cognate tRNA in the cytosol (Fu et al., 2012). Consistently, we found that angiogenin, a stress-activated ribonuclease that cleaves tRNA at the anticodon loop (Li and Hu, 2012; Thompson and Parker, 2009), stimulates the nuclear translocation of TyrRS (Wei et al., 2014). In fact, TyrRS nuclear translocation can be stimulated by a wide variety of stress conditions, including oxidative stress, ER stress, heat shock, and serum starvation (Sajish and Schimmel, 2015; Wei et al., 2014). The nucleus-localized TyrRS strongly protects against UV-induced DNA double-strand breaks in zebrafish and does so by upregulating DNA damage repair genes, such as BRCA1 and RAD51, through activating transcription factor E2F1 (Wei et al., 2014). In consequence, TyrRS has been established as an important stress-response protein through its nuclear functions. However, the role of nuclear TyrRS in translation control has not been investigated so far. Due to the established knowledge on tRNA regulation, we speculated that TyrRS might also plays an important role in stress-induced adaptive translation regulation through cytosol-nucleus translocation.

We set out to investigate this hypothesis and found that nuclear TyrRS inhibits global translation by binding to and suppressing genes crucial to protein synthesis. Importantly, the effect of nuclear TyrRS in translation control happens after the mTOR and eIF2α responses and thus provides the cells with a second chance to adapt and survive through the stress conditions. Apart from this difference in timing, the ability of TyrRS to respond to a wide variety of stress factors and to effect translation inhibition while selectively activating stress-response genes resembles the function of ISR. Therefore, TyrRS may be considered as the core effector for an ISR-like response to aid cells during prolonged stress. This late integrated stress response may also involve other tRNA synthetase family members to collectively sense a larger variety of stress factors and instigate diverse, protective responses to restore cellular homeostasis through their cytosol-nucleus translocations.

## Results

### Nuclear TyrRS inhibits global translation during late stage oxidative stress

Oxidative stress is able to activate all kinases upstream of eIF2α (Harding et al., 2003; Pyo et al., 2008; Suragani et al., 2012; Baker et al., 2012; Pakos-Zebrucka et al., 2016) as well as to inhibit mTOR signaling (Kim et al., 2002; Reiling and Sabatini, 2006) in mammalian cells. Therefore, we set out to evaluate the effect of TyrRS nuclear entry on global translation under oxidative stress to allow for direct comparison with the eIF2α and/or mTOR regulated stress response. We took advantage of our previously developed approach to specifically exclude TyrRS from the nucleus without affecting its cytoplasmic role in aminoacylation (Wei et al., 2014). Briefly, we knocked down endogenous TyrRS in HEK293 cells and compensated by expressing a wild-type or NLS-mutated form of TyrRS, to generate ΔY/YARS (“normal”) or ΔY/YARS-NLS^Mut^ (nuclear TyrRS deficient) stable cell lines, respectively. We confirmed that the expression level of ectopically expressed TyrRS in both cells is similar to that of endogenous TyrRS (Supplementary Fig. S1). We have also confirmed that H_2_O_2_ treatment, which induces oxidative stress, stimulates significant amounts of TyrRS to enter the nucleus in ΔY/YARS cells as in HEK293 cells, but the amount of nuclear TyrRS is substantially reduced in ΔY/YARS-NLS^Mut^ (Supplementary Fig. S1). With this system in place, we here examined the cellular level of global protein synthesis using the SUrface Sensing of Translation (SUnSET) technique (Schmidt et al., 2009). As shown in Fig. 1A and quantified in Fig. 1B, H_2_O_2_ treatment inhibits global translation in a time-dependent manner. Shortly after H_2_O_2_ treatment (1h), a sharp decrease (~50%) in global protein synthesis was observed in both “normal” and nuclear TyrRS deficient HEK293 cells. However, after a longer period of H_2_O_2_ treatment (4h, 12h, or 24h), the global translation level in nuclear TyrRS deficient cells was strikingly different from that in “normal” cells. In “normal” cells, the global translation level recovered to about 70% level after 4h and this level was maintained during further H_2_O_2_ treatment; however, nuclear TyrRS deficient cells exhibit faster recovery of translation and also reached a higher than normal level of translation after 12h of H_2_O_2_ treatment. These results suggest that nuclear TyrRS functions to inhibit and prevent overstimulation of global translation when cells are still under oxidative stress.

**Figure 1.**
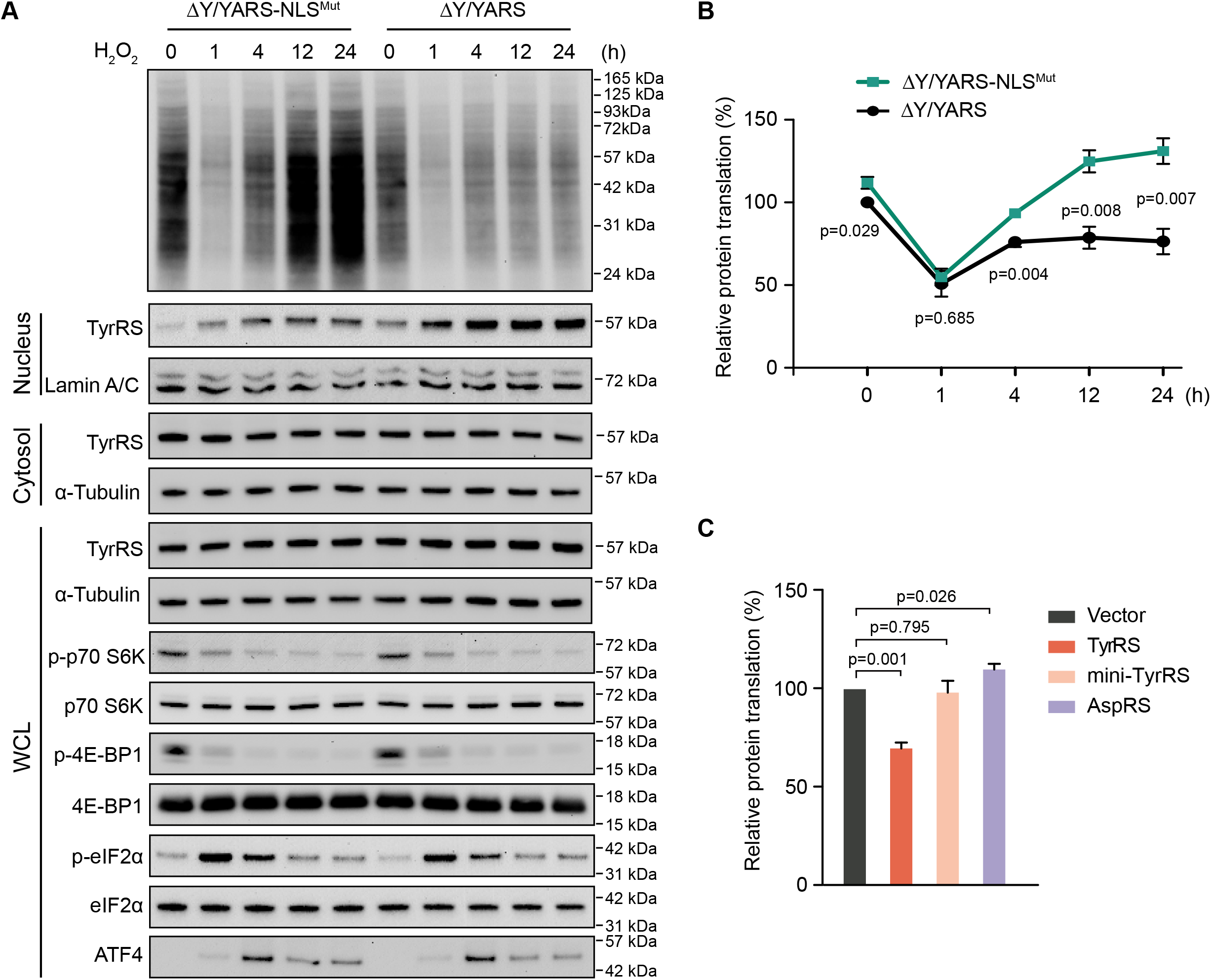
Nuclear TyrRS inhibits global protein translation during late stage oxidative stress. A) SUnSET analysis to detect cellular protein synthesis activity in HEK293 cells with or without nuclear TyrRS deficiency. Cells were treated with H_2_O_2_ for the indicated time. The nuclear localization of TyrRS was analyzed by cell fractionation and then Western blot. Total TyrRS expression levels and markers for the eIF2a and mTOR signaling pathway were analyzed by Western blot using the whole cell lysate (WCL). ΔY/YARS-NLS^Mut^: HEK293 cells with a knock down of endogenous TyrRS and expression of TyrRS with a mutated NLS (^242^KKKLKK^247^ to ^242^<u>NN</u>KL<u>N</u>K^247^). ΔY/YARS: HEK293 cells with a knock down of endogenous TyrRS and ectopic expression of wild type TyrRS. Lamin A/C: nuclear marker, α-Tubulin: cytoplasmic marker. B) Quantification of protein synthesis activity measured by SUnSET. The intensities of the signal indicating newly synthesized proteins were normalized to α-Tubulin for each treatment. Normalized protein synthesis activity in ΔY/YARS at time point 0 was set at 100%. n=3, biological replicates, one way Student’s t test. C) Quantified SUnSET analysis to show protein synthesis inhibition by overexpression of wild type TyrRS. Overexpression of mini-TyrRS or AspRS showed no inhibition of translation. n=3, biological replicates, one way Student’s t test. Equal expression of the various transgenes (TyrRS, mini-TyrRS, or AspRS) in HEK293 cells were confirmed by Western Blot analysis with the loading control (α-tubulin) and representatively shown in Supplementary Fig. 2.

### The mTOR and eIF2α pathways do not contribute to the inhibitory effect of nuclear TyrRS

Next, we studied the nuclear TyrRS-mediated global translation inhibition in relationship to mTOR and eIF2α pathways. The mTOR pathway is one of the most well studied translational control mechanisms during oxidative stress (Kim et al., 2002; Reiling and Sabatini, 2006). Many amino acids and other types of nutrients can lead to the activation of mTOR (Bar-Peled and Sabatini, 2014). mTOR promotes protein synthesis by phosphorylating the eukaryotic initiation factor 4E (eIF4E)-binding proteins (4E-BPs) and the p70 ribosomal S6 kinase (S6K) (Reiling and Sabatini, 2006; Spriggs et al., 2010). The phosphorylation of 4E-BPs prevents its binding to eIF4E, enabling eIF4E to promote cap-dependent translation (Marcotrigiano et al., 1999). The phosphorylation of S6K leads to its activation and subsequent stimulation of ribosomal protein synthesis (Ruvinsky et al., 2005). We assessed mTOR activity in ΔY/YARS and ΔY/YARS-NLS^Mut^ HEK293 cells by measuring the phosphorylation of its substrates 4E-BP1 and S6K1. As shown in Fig. 1A, the phosphorylation levels of 4E-BP1 and S6K1 decreased shortly after H_2_O_2_ treatment (1h), and this effect was sustained during the 24h H_2_O_2_ treatment. Nevertheless, there was no obvious difference in the phosphorylation of 4E-BP1 and S6K1 between ΔY/YARS and ΔY/YARS-NLS^Mut^ cells, suggesting nuclear TyrRS inhibits protein translation through an mTOR-independent mechanism.

The ISR was assessed through its core regulators eIF2α and ATF4. We analyzed the phosphorylation of eIF2α and the expression of ATF4 in ΔY/YARS and ΔY/Y ARS-NLS^Mut^ HEK293 cells under oxidative stress. Consistent with a previous report (Shenton et al., 2006), phosphorylation of Ser^51^ in eIF2α and the subsequent expression of ATF4 were induced shortly after H_2_O_2_ treatment (Fig. 1A). However, there was no obvious difference in eIF2α phosphorylation and ATF4 expression between ΔY/YARS and ΔY/YARS-NLS^Mut^ cells at any of the tested time points, suggesting nuclear TyrRS inhibits protein translation independent of ISR activation.

Importantly, both the reduction of mTOR activity and the phosphorylation of eIF2α happened quickly after H_2_O_2_ treatment, suggesting that they were responsible for the initial decrease in global protein synthesis in both “normal” and nuclear TyrRS deficient HEK293 cells (Fig. 1A). In contrast, the effect of nuclear TyrRS on translation manifested at a later time point, suggesting a slower, possibly transcriptional mechanism.

To exclude the possibility that oxidative stress-induced nuclear translocation of TyrRS inhibits protein synthesis simply by depleting the cytosolic pool of the synthetase, we increased TyrRS availability by overexpressing V5-tagged wild-type TyrRS in HEK293 cells (Supplementary Fig. S2). Interestingly, overexpression of TyrRS did not promote but rather suppressed global translation (Fig. 1C). As overexpression of TyrRS would also increase its level in the nucleus (Wei et al., 2014), this observation is consistent with a nuclear mechanism underlying the translation inhibitory effect of TyrRS.

### Nuclear TyrRS binds to DNA elements in translation-related genes

Given the binding ability of TyrRS to nucleic acids in the cytoplasm (tRNA), we speculated that TyrRS may be able to bind certain DNAs in the nucleus, for example of genes that are relevant for translation.

We generated a V5-tagged-TyrRS-expressing stable HEK293 cell line and performed a chromatin immunoprecipitation (ChIP)-sequencing experiment by utilizing an α-V5 antibody. A stable HEK293 cell line with only V5-tag expression was used as a control. We found a total of 20 DNA sites that were bound by TyrRS-V5 (Supplementary Table S1). The vast majority of these binding sites (16/20) fall within regions of protein-coding genes. Remarkably, 6 out of these genes encode translation-related proteins, including TyrRS itself (YARS) and three other tRNA-synthetases (SARS, WARS, and GARS), eukaryotic elongation factor (EEF1A1), and mRNA nuclear export gene (RAE1) (Fig. 2A).

**Figure 2.**
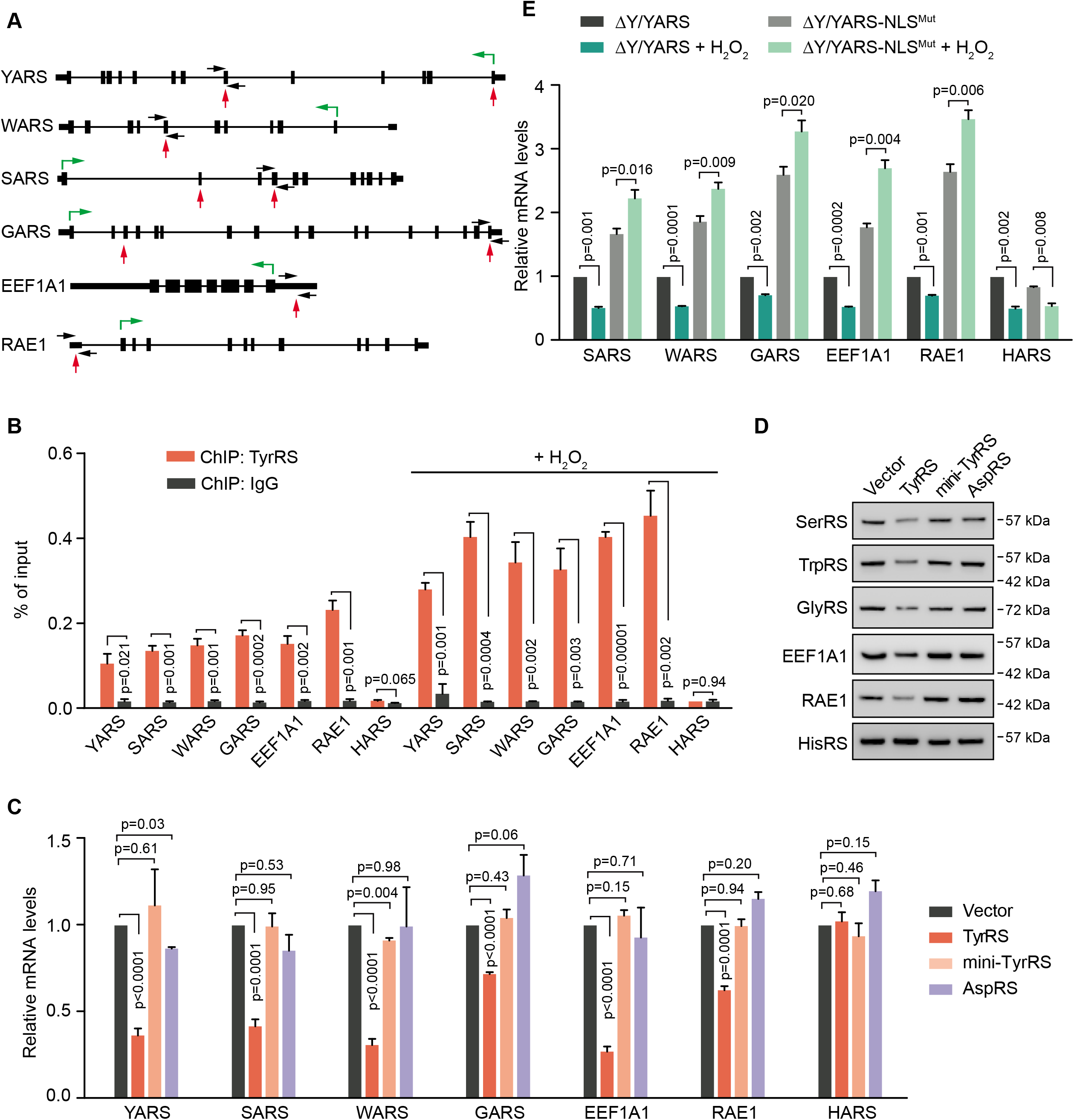
Nuclear TyrRS binds to DNA elements of translation-related genes and represses their transcription. A) Illustration of TyrRS binding sites on its target genes that are related to protein translation as identified by ChIP-seq. Black boxes, exons. Black thick lines, 5’ UTR or 3’ UTR. Black thin lines, introns. Red arrows, TyrRS binding sites. Black arrows, primers for ChIP-qPCR. Green arrows, directions and starting sites of translation. B) ChIP-qPCR confirms the DNA binding ability of endogenous TyrRS on protein translation-related genes. H_2_O_2_ treatment for 12 hours enhances TyrRS DNA binding ability. n=3, biological replicates, one way Student’s t test. C) Overexpression of TyrRS in HEK293 cells represses transcription of translation-related genes. Overexpression of mini-TyrRS or AspRS shows no repression of TyrRS target gene transcription. Transcription of target genes were measured by RT-PCR using HEK293 cells with transgene overexpression for 24 hours. n=3, biological replicates, one way Student’s t test. Equal expression of the various transgenes (TyrRS, mini-TyrRS, or AspRS) in HEK293 cells were confirmed by Western Blot analysis with the loading control (α-tubulin) and representatively shown in Supplementary Fig. 2. D) Overexpression of full length TyrRS, but not mini-TyrRS or AspRS, downregulates its target genes on protein level 48 hours after transfection. Equal expression of the various transgenes (TyrRS, mini-TyrRS, or AspRS) in HEK293 cells were confirmed by Western Blot analysis with the loading control (α-tubulin) and representatively shown in Supplementary Fig. 2. E) Restricting TyrRS nuclear localization in HEK293 cells promotes transcription of protein translation-related genes. H_2_O_2_ treatment for 12 hours inhibits transcription of TyrRS target genes in normal cells, but not in TyrRS nuclear location deficient cells. n=3, biological replicates, one way Student’s t test. ΔY/YARS-NLS^Mut^: HEK293 cells with a knock down of endogenous TyrRS and expression of TyrRS with a mutated NLS (^242^KKKLKK^247^ to ^242^<u>NN</u>KL<u>N</u>K^247^). ΔY/YARS: HEK293 cells with a knock down of endogenous TyrRS and ectopic expression of wild type TyrRS.

To confirm TyrRS binding at these translation-related genes, we performed a ChIP-PCR assay. A tRNA synthetase gene not bound by TyrRS in our ChIP-sequencing experiment (i.e., HARS) was used as a negative control. Indeed, the TyrRS-V5 protein overexpressed in HEK293 cells significantly enriched genetic elements from YARS, SARS, WARS, GARS, EEF1A1, and RAE1, but not HARS (Supplementary Fig. S3). We also confirmed the binding of endogenous TyrRS from untransfected HEK293 cells to these target sites (Fig. 2B). Because oxidative stress promotes the nuclear translocation of TyrRS (Wei et al., 2014) (Supplementary Fig. S1), the binding of TyrRS to its target sites was greatly enhanced under H_2_O_2_ treatment as expected (Fig. 2B). These results strongly suggest that TyrRS may regulate the expression of these translation-related genes to inhibit global translation in response to oxidative stress.

### Nuclear TyrRS inhibits transcription of translation-related genes

To test the effect of TyrRS on transcription of its target genes, we again ectopically expressed TyrRS in HEK293 cells (Supplementary Fig. S2). To understand if the potential effect is unique to TyrRS, we separately expressed another tRNA synthetases as a control - aspartyl-tRNA synthetase (AspRS or DARS) (Supplementary Fig. S2). We also tested mini-TyrRS, the catalytic core of TyrRS that is constituted of the catalytic domain and the anticodon binding domain, where the NLS is located (Wakasugi and Schimmel, 1999). Mini-TyrRS is the evolutionarily conserved region across all forms of life, which is linked to a C-terminal EMAP II-like domain (CTD) that only appeares in TyrRS in vertebrates and insects and is dispensable for enzymatic activity (Wakasugi and Schimmel, 1999). Real-time PCR results showed that the transcript levels of TyrRS target genes, but not of the control gene HARS, were uniformly inhibited by TyrRS overexpression (Fig. 2C). In contrast, overexpression of AspRS or mini-TyrRS did not significantly affect the transcript level of TyrRS target genes (Fig. 2C and Supplementary Fig. S2). The changes in protein level of these genes as detected by Western blot analysis were consistent with the changes found on transcript level (Fig. 2D and Supplementary Fig. S2). These changes were also consistent with our observation that overexpression of TyrRS, but not mini-TyrRS or AspRS, in HEK293 cells repressed global protein translation (Fig. 1C), suggesting TyrRS inhibits global translation through transcriptionally repressing certain translation-related genes.

To confirm that the repressive effect of TyrRS was not due to its overexpression but due to its nuclear localization upon oxidative stress, we also tested the transcription level of TyrRS target genes in ΔY/YARS and ΔY/YARS-NLS^Mut^ HEK293 cells. Excluding TyrRS from the nucleus greatly increased the transcription of SARS, WARS, GARS, EEF1A1, and RAE1, but not HARS (Fig. 2E), demonstrating that nuclear TyrRS at endogenous expression levels is able to repress the transcription of its target genes. Interestingly, in “normal” (ΔY/YARS) cells, 12 h of H_2_O_2_ treatment significantly decreased the transcript level of TyrRS target genes. In contrast, the transcript levels of TyrRS target genes was significantly increased, rather than decreased, in nuclear TyrRS deficient (ΔY/YARS-NLS^Mut^) cells under H_2_O_2_ treatment (Fig. 2E). TyrRS non-target gene HARS transcription was decreased in both “normal” and nuclear TyrRS deficient cells after 12h H_2_O_2_ treatment (Fig. 2E), suggesting that HARS transcription was also suppressed under oxidative stress but through a mechanism independent of nuclear TyrRS. These results correlate well with our earlier observation that nuclear TyrRS deficient cells exhibit a higher than normal level of translation after 12h under H_2_O_2_ treatment (Fig. 1A). It further strengthens the conclusion that nuclear TyrRS inhibits global translation at later stages of oxidative stress through a transcriptional mechanism to repress the expression of translation-related genes.

### Nuclear TyrRS recruits TRIM28/HDAC1 or NuRD complex to its target genes

Active genomic regulatory elements are often flanked by histones with H3 acetylation modification at lysine 27 residue (H3K27Ac) (Shlyueva et al., 2014). To understand how TyrRS represses its target genes’ transcription, we looked into histone H3K27Ac modifications on TyrRS binding sites by ChIP assay. As shown in Fig. 3A, overexpressing TyrRS in HEK293 cells remarkably reduced the H3K27Ac levels on TyrRS target genes, without a significant effect on the control HARS gene. This result suggests that TyrRS may recruit some histone modifiers to epigenetically regulate expression of its target genes.

**Figure 3.**
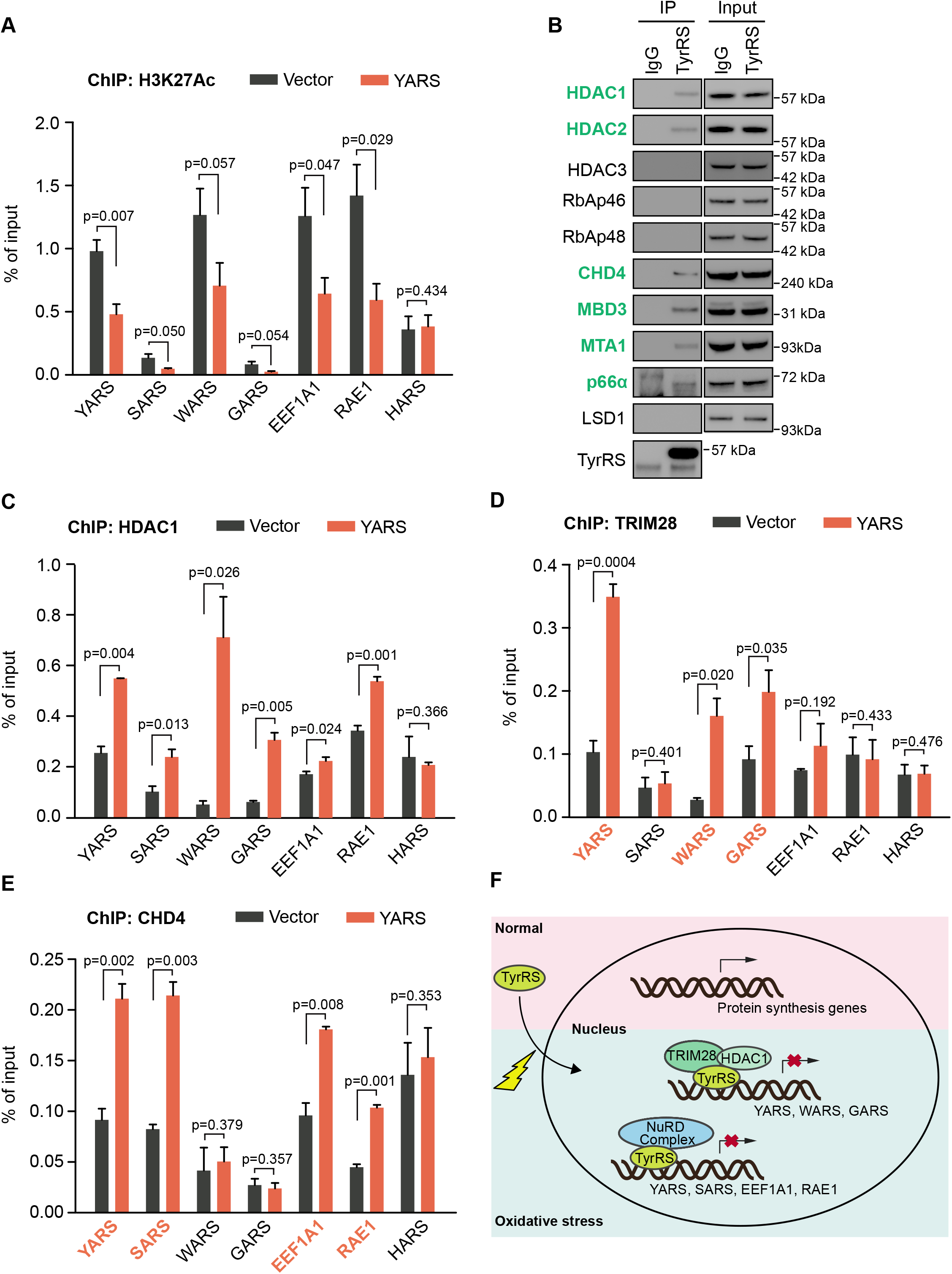
TyrRS recruits transcriptional co-repressor TRIM28/HDAC1 or NuRD complex to epigenetically repress expression of its target genes. All ChIP-qPCR assays were done using HEK293 cells with vector or TyrRS overexpression for 24 hours. A) TyrRS represses histone H3 acetylation on its target sites. The enrichment of histone H3 acetylation was determined by chromatin IP using α-H3K27Ac antibody followed by qPCR. n=3, biological replicates, one way Student’s t test. Vector: control, YARS: TyrRS overexpression. B) TyrRS interacts with HDAC1 and other factors in the NuRD complex. Co-immunoprecipitation using a-TyrRS antibody followed by Western blot analysis to detect proteins involved in the NuRD complex. Vector: control, YARS: TyrRS overexpression. C) TyrRS recruits HDAC1 to its target sites on protein translation-related genes in TyrRS overexpressing cells. Enrichment of HDAC1 was detected by chromatin IP using α-HDAC1 antibody followed by qPCR. n=3, biological replicates, one way Student’s t test. Vector: control, YARS: TyrRS overexpression. D) Recruitment of TRIM28 at multiple TyrRS target sites (YARS, WARS, and GARS) in TyrRS overexpressing HEK293 cells. Enrichment of TRIM28 was detected by chromatin IP using α-TRIM28 antibody followed by qPCR. n=3, biological replicates, one way Student’s t test. Vector: control, YARS: TyrRS overexpression. E) Increased occupancy of CHD4, a component of the NuRD complex factor, at multiple TyrRS target sites (YARS, SARS, EEF1A1, and RAE1) in TyrRS overexpressing HEK293 cells. Enrichment of CHD4 was detected by chromatin IP using α-CHD4 antibody followed by qPCR. n=3, biological replicates, one way Student’s t test. Vector: control, YARS: TyrRS overexpression. F) Schematic illustration of how TyrRS responds to oxidative stress to repress transcription of protein translation-related genes.

Several histone deacetylases, including HDAC1, HDAC2, and HDAC3, were identified to interact with TyrRS in HeLa cells (Wei et al., 2014). Interestingly, we found that the interaction between TyrRS and HDAC1 is not direct, but mediated by transcription co-repressor TRIM28 (Wei et al., 2014). To confirm these interactions and to identify other potential interaction partners of TyrRS in HEK293 cells, we performed a stringent Tandem Affinity Purification and Mass Spectrometry (TAP-MS) analysis. TyrRS was fused with two small, but high affinity tags, a streptavidin binding peptide (SBP) and a calmodulin binding peptide (CBP), to enable two successive purification steps to enhance the specificity of identified interaction partners (Supplementary Fig. S4). The potential interaction partners of TyrRS identified by this method are listed in Supplementary Table S2. We confirmed HDAC2 and TRIM28 as TyrRS interactors in HEK293 cells, and also identified two other histone modification related proteins, CHD4 and MBD3 as candidate interaction partners of TyrRS (Supplementary Table S2). Intriguingly, CHD4, MBD3, HDAC2, and HDAC1 are all components of the Nucleosome Remodeling Deacetylase (NuRD) complex, a major chromatin remodeling complex, which also contains other components, including RbAp46, RbAp48, CHD4, MBD3, MTA1, p66α, and LSD1.

We next confirmed the interaction between endogenous TyrRS and these factors found by our interactome study with Co-IP. Indeed, TyrRS can interact with HDAC1, HDAC2, CHD4, and MBD3 in HEK293 cells (Fig. 3B). In addition, we could retrieve two other factors of the NuRD complex, which were not found in the initial interactome study, MTA1 and p66α (Fig. 3B). These results highly suggest that in the nucleus, TyrRS can recruit different co-repressors (TRIM28/HDAC1 or NuRD complex) to epigenetically regulate target genes transcription.

To test this hypothesis, we evaluated the effect of TyrRS on binding of these transcriptional corepressors to the TyrRS target genes in HEK293 cells by ChIP assay. We first tested HDAC1, which is a shared co-factor for both TRIM28 and the NuRD complex. Over-expression of TyrRS significantly increased the amount of HDAC1 bound to all translation-related TyrRS target genes (Fig. 3C). In contrast, TRIM28 was recruited by TyrRS to 3 of its target genes (YARS, WARS, and GARS) (Fig. 3D), while CHD4 was recruited to a different gene set (YARS, SARS, EEF1A1, and RAE1) (Fig. 3E). YARS was the only gene bound by both the NuRD complex (as represented by CHD4) and TRIM28. Importantly, all translation-related TyrRS target genes were bound by either the NuRD complex or TRIM28, or both.

To strengthen the evidence that these co-factors are required for TyrRS to regulate target gene expression and because HDAC1 is a shared co-factor for both TRIM28 and the NuRD complex, we investigated whether an HDAC1 inhibitor could abolish the transcriptional repressor activity of TyrRS. Indeed, in the presence of Trichostatin A (TSA), a selective inhibitor of the class I and II HDACs (including HDAC1) (Schultz et al., 2004; Bieliauskas and Pflum, 2008), overexpression of TyrRS no longer inhibited the expression of its target genes (Supplementary Fig. S5).

### Gain-of-function CMT-causing mutation E196K over-represses translation

TyrRS is one of the several tRNA synthetases causatively linked to a hereditary neurodegenerative condition called Charcot-Marie-Tooth (CMT) disease (Jordanova et al., 2006). Using a *Drosophila* model, we recently linked the nuclear localization of TyrRS to CMT disease (Bervoets et al., 2019). We also showed that CMT-causing TyrRS mutants, including E196K TyrRS, are gain-of-function and undergo stronger interactions with HDAC1 in the nucleus (Bervoets et al., 2019; Blocquel et al., 2017). If our proposed mechanism (i.e., nuclear TyrRS inhibits global translation through recruiting HDAC1 complexes to translation related genes to epigenetically repress their expression) as illustrated in Fig. 3F is correct, we would expect a CMT-causing mutant TyrRS protein to over-inhibit the transcription of TyrRS target genes and global translation. Indeed, compared to wild-type TyrRS, E196K TyrRS significantly enhanced the inhibitory effect of TyrRS on global translation (Supplementary Fig. S6A and S6B) and specifically on transcription of most of its target genes (Supplementary Fig. S6C).

### Nuclear TyrRS is protective against cell death induced by oxidative stress

Dysregulation of protein synthesis often leads to cell death (Anderson et al., 2009; Boyce et al., 2005; Tsaytler et al., 2011). Therefore, we evaluated the effect of nuclear TyrRS on cell viability under oxidative stress. After 12 h of H_2_O_2_ treatment, the nuclear TyrRS deficient (ΔY/YARS-NLS^Mut^) cells exhibited more cell death and reduced resistance to oxidative stress than “normal” (ΔY/YARS) cells (Fig. 4A). The reduced viability of ΔY/YARS-NLS^Mut^ cells was accompanied by increased levels of cleaved caspase 3 (Fig. 4B and Supplementary Fig. S7), suggesting that oxidative stress enhanced the activation of a cell death program in nuclear TyrRS deficient cells. In consequence, nuclear TyrRS is protective against cell death induced by oxidative stress. Importantly, this effect of nuclear TyrRS was independent, at least in part, of its protective role against DNA damage, which is mediated by transcriptional activation of E2F1 (Wei et al., 2014): Knocking down E2F1 reduced the viability of both “normal” and nuclear TyrRS deficient cells (Supplementary Fig. S8A and S8B), however, the cell protective effect of nuclear TyrRS against oxidative stress was largely maintained, as despite the E2F1 knockdown, ΔY/YARS cells showed stronger resistance to oxidative stress than ΔY/YARS-NLS^Mut^ cells (Supplementary Fig. S8A and S8B).

**Figure 4.**
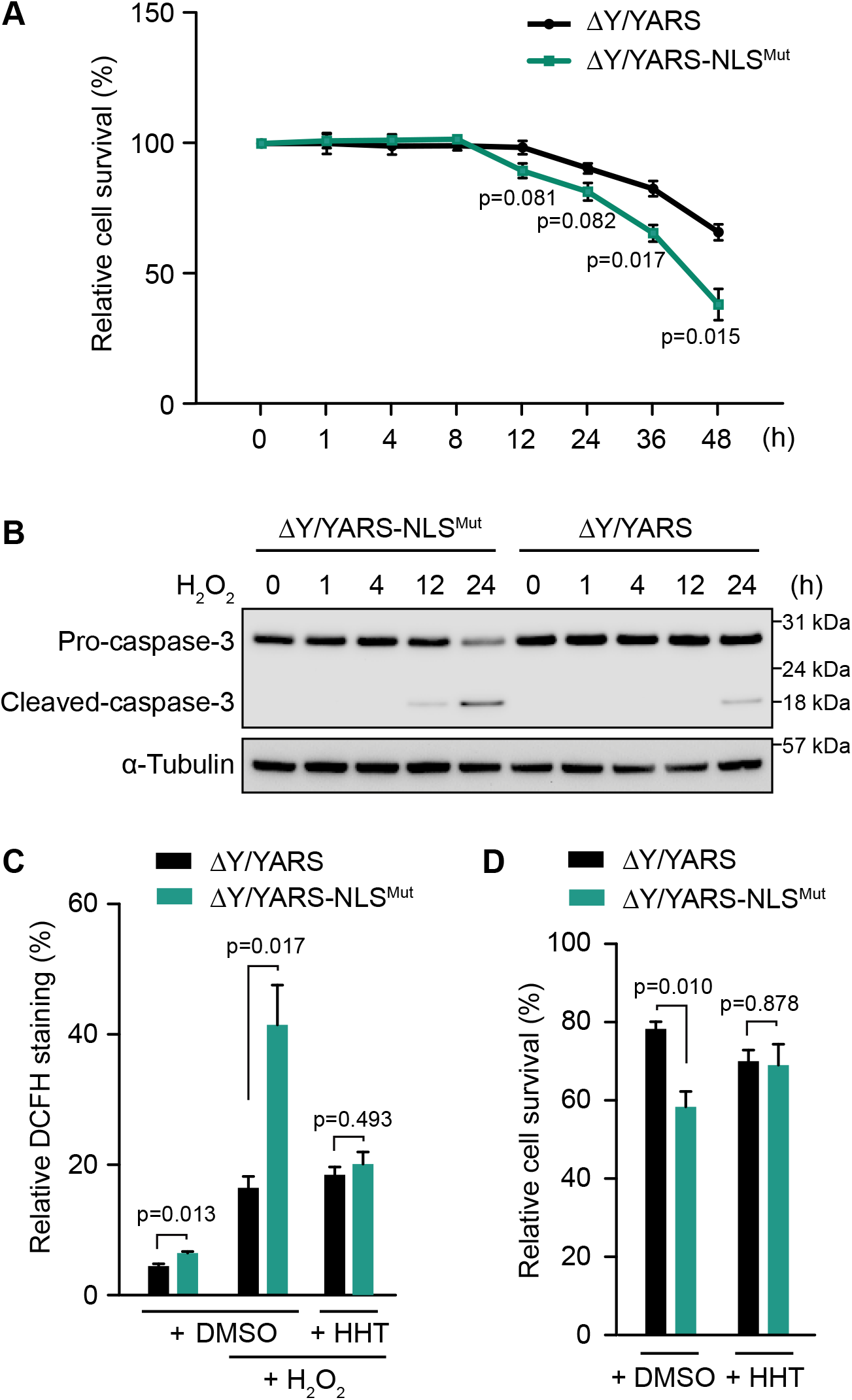
Nuclear TyrRS protects against cell death induced by oxidative stress. ΔY/YARS-NLS^Mut^: HEK293 cells with a knock down of endogenous TyrRS and expression of TyrRS with a mutated NLS (^242^KKKLKK^247^ to ^242^<u>NN</u>KL<u>N</u>K^247^). ΔY/YARS: HEK293 cells with a knock down of endogenous TyrRS and ectopic expression of wild type TyrRS. A) Exclusion of TyrRS from the nucleus enhances cell death caused by H_2_O_2_ treatment. Relative cell viabilities were measured with a cell counting kit. Cell survival in ΔY/YARS at time point 0 was set to 100%. n=3, biological replicates, one way Student’s t test. B) Enhanced apoptosis in nuclear TyrRS deficient HEK293 cells. Caspase-3 cleavage, a marker of apoptosis, was detected by Western blot analysis. C) Nuclear TyrRS prevents ROS over-accumulation under H_2_O_2_ treatment for 36 hours. ROS accumulation was detected by the general oxidative stress indicator CM-H_2_DCFDA and shown as relative DCFH staining. n=3, biological replicates, one way Student’s t test. D) Inhibition of protein synthesis by homoharringtonine (HHT) treatment reduces cell death in nuclear TyrRS deficient cells. Cells were treated with or without HHT (50 mM) for 24 hours and followed by H_2_O_2_ treatment for 36 hours. Relative cell viabilities were measured with a cell counting kit and normalized to viability of ΔY/YARS seeded at the same time but without HHT and H_2_O_2_ treatment. n=3, biological replicates, one way Student’s t test.

### Cell protection by nuclear TyrRS is predominately mediated by its impact on translation

Stress-induced cell death has been linked to overactivation of translation and subsequent intracellular ROS accumulation (Han et al., 2013; Marciniak et al., 2004). To understand if nuclear TyrRS also affects ROS accumulation, we measured ROS by CM-H_2_DCFDA staining in ΔY/YARS and ΔY/YARS-NLS^Mut^ cells (Fig. 4C). Interestingly, even without H_2_O_2_ treatment, the ROS levels in nuclear TyrRS deficient cells was higher than in “normal” cells. During H_2_O_2_ treatment, the difference in ROS levels in ΔY/YARS and ΔY/YARS-NLS^Mut^ cells was dramatically increased (Fig. 4C). However, this difference in ROS levels was abolished if we pre-treated the cells with the translation elongation inhibitor homoharringtonine (HTT) (Fig. 4C), suggesting that increased intracellular levels of ROS in nuclear TyrRS deficient cells were translation dependent. Consistent with this conclusion, no obvious difference in the levels of antioxidative stress response genes, such as SOD1, catalase, and Ucp2 was observed in ΔY/YARS versus ΔY/YARS-NLS^Mut^ cells (Supplementary Fig. S9). This observation rules out the possibility that nuclear TyrRS reduced intracellular ROS accumulation by promoting an anti-oxidative stress response. Interestingly, HTT treatment also completely rescued the loss of viability in nuclear TyrRS deficient cells compared to “normal” cells (Fig. 4D), confirming that the protective effect of nuclear TyrRS against cell death is predominantly mediated by its impact on translation.

## Discussion

Through their evolutionarily conserved enzymatic function in charging tRNA with their cognate amino acid, aaRSs are essential for protein synthesis in all domains of life. In complex multicellular organisms, the functional landscape of aaRSs is expanded with broad regulatory functions (Guo et al., 2010; Guo and Schimmel, 2013; Kwon et al., 2019). However, the connections between their classical enzymatic role and their non-enzymatic functions remain to be understood. TyrRS was the first aaRS shown to have regulatory roles, among them several in the nucleus (Wakasugi and Schimmel, 1999; Wei et al., 2014; Sajish and Schimmel, 2015). When the NLS of TyrRS was identified, its special location immediately indicated a control of the nuclear import of the synthetase by its tRNA, which was subsequently demonstrated (Fu et al., 2012). This regulation and the increasing evidence for the involvement of tRNA in translational control during stress response (Dong et al., 2000; Tahmasebi et al., 2018) led us to speculate a direct role of TyrRS nucleus translocation in stress response as well. Indeed, we found that the nuclear import of TyrRS is stimulated under stress and that nucleus-localized TyrRS functions through the transcriptional machinery to promote the expression of DNA damage response genes for cell protection(Wei et al., 2014). Here in this work, we reveal that nuclear TyrRS also inhibits global translation by binding to protein synthesis related genes and recruiting TRIM28/HDAC1 or the NuRD complex to repress their transcription.

It has been demonstrated that a variety of stressors including UV exposure, heat stress, serum starvation, H_2_O_2_, sodium arsenite, and tunicamycin treatment can trigger TyrRS nucleus translocation (Sajish and Schimmel, 2015; Wei et al., 2014). We propose that these stress signals are transmitted through intrinsic properties of the tRNA synthetase enzyme. The aminoacylation reaction happens in two steps: In the first step, TyrRS binds to its amino acid and ATP to catalyze the formation of an enzyme-bound tyrosyl-adenylate, which triggers tRNA binding for the second step: the activated amino acid subsequently reacts with tRNA to yield aminoacyl-tRNA (Fig. 5). For most aaRSs, including TyrRS, the second step or the release of the aminoacyl-tRNA product is rate-limiting (Zhang et al., 2006). Therefore, in unstressed cells, most TyrRS in the cytosol exists in a tRNA-bound state in which the NLS is blocked to prevent its nuclear translocation (Fig. 5). Depletion of tRNA leads to TyrRS NLS exposure and nuclear translocation as we demonstrated (Fu et al., 2012). Under a variety of stress conditions, including oxidative stress, heat stress, UV exposure, serum starvation, cold stress, and hypoxia, tRNAs are cleaved by angiogenin (Ivanov et al., 2011; Thompson and Parker, 2009). Depletion of tRNA can also result from retrograde nuclear tRNA import, which is also increased upon oxidative stress (Schwenzer et al., 2019). Specifically tRNA^Tyr^ is fragmented in response to oxidative stress, resulting in lower levels of mature tRNA (Huh et al., 2018). NLS exposure can also be achieved by modification of the synthetase, independent of the status of tRNA. A recent report showed that enhanced TyrRS nuclear localization during oxidative stress is mediated through lysine acetylation at its NLS (Cao et al., 2017), which is likely to weaken tRNA binding, thereby constantly exposing the NLS. Another mechanism to promote TyrRS nuclear translocation may be through depleting the substrates (i.e., ATP and amino acid), which would slow down the first step of the aminoacylation reaction, when the synthetase is free from tRNA binding (Fig. 5). Indeed, inhibiting the enzymatic activity of TyrRS through resveratrol, a tyrosine-like molecule, was shown to stimulate the nuclear translocation of the synthetase (Sajish and Schimmel, 2015). Therefore, through its active site TyrRS has the intrinsic capability to sense amino acid and energy deficiency within the cell. Importantly, this type of metabolic stress can be integrated with other tRNA-mediated stress stimuli through the nuclear translocation of TyrRS, which subsequently leads to global translation inhibition and stress-response genes activation.

**Figure 5.**
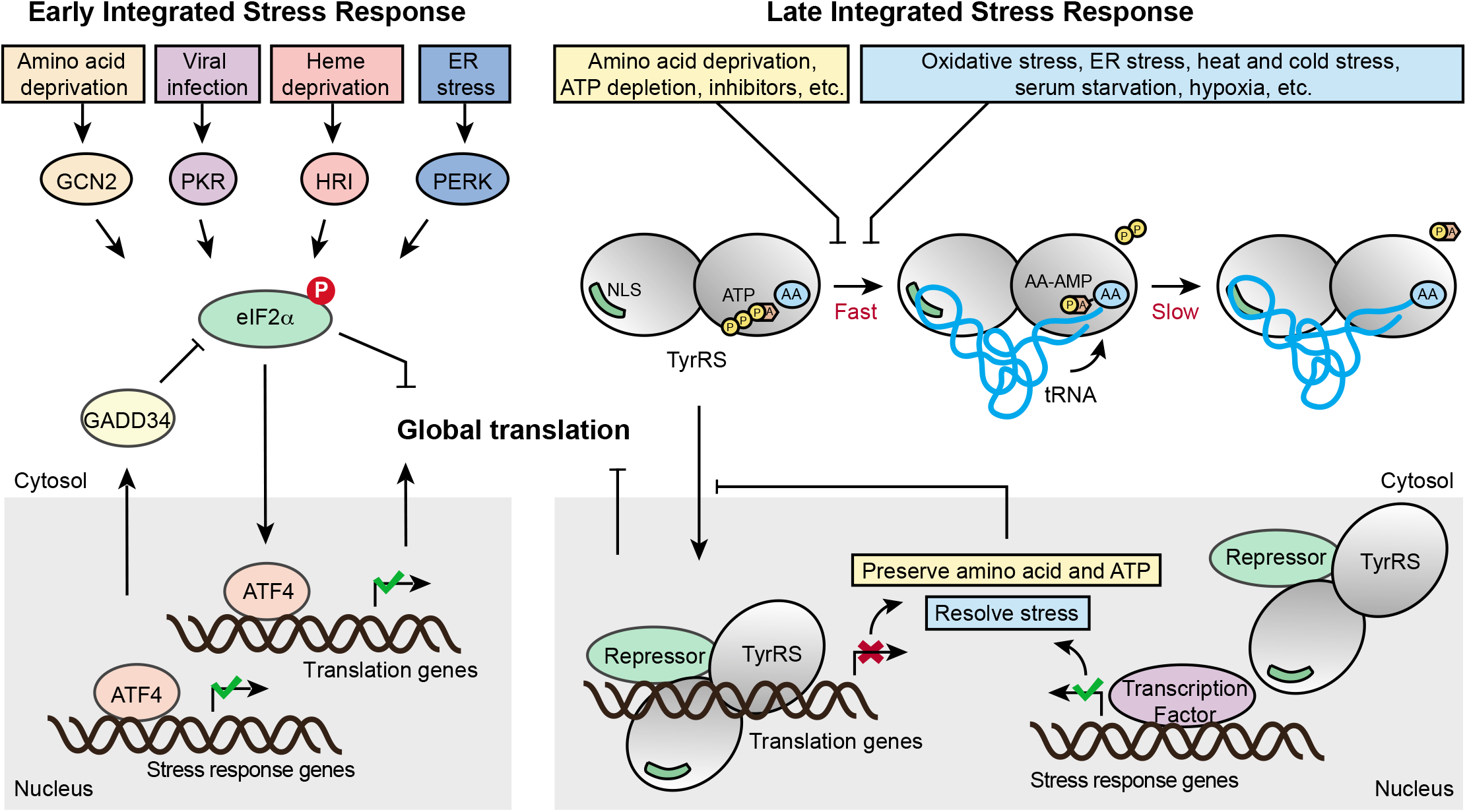
Schematic illustration of the classical ISR and the TyrRS-mediated late ISR pathways in parallel. As an immediate response to a wide range of stress signals, the classical ISR pathway is activated by four upstream kinases that phosphorylate eIF2α to subsequently inhibit global translation while selectively activating transcription factor ATF4 to aid cell survival and recovery. While the classical ISR provides the early response, the tRNA synthetase-based ISR mediates the late response, providing the cells with a second chance to adapt and survive through stress conditions. A variety of stress conditions can be sensed by TyrRS through its intrinsic properties as a tRNA synthetase and its strategically located NLS and be integrated through TyrRS nuclear translocation to effect protective responses against prolonged stress. Similar to the negative feedback regulation instigated in ISR to prevent over-suppression of translation and to restore homeostasis, the tRNA synthetase-mediated response also provides negative feedback mechanisms: The translation inhibition and stress-response genes activation effected by nuclear TyrRS would preserve ATP and amino acid and help alleviate the stressors that stimulate TyrRS nuclear translocation in the first place.

The manner of activation of TyrRS described here is reminiscent of the activation of another key regulator of cellular response, the hypoxia-inducible factor HIF-1. HIF-1 is actively kept in an inactive state by constant degradation of one of its subunits, HIF-1α (Semenza, 2001). In direct response to a lack of oxygen or other inducers HIF-1α degradation stops and HIF-1 can act (Semenza, 2001). This failsafe mechanism eschews the need for an active step to initiate its transcription factor function which could fail if resources are limited. In an analogous manner, the TyrRS-dependent integrated stress response is activated by the depletion of its negative regulators of nuclear localization, tRNA^Tyr^, ATP, and tyrosine. The lack of an active on-switch together with the ability to intrinsically sense small molecules and tRNA status allows TyrRS to act autonomously from other cell signaling molecules, only connected by metabolic feedback.

Similar to the negative feedback regulation instigated in ISR to prevent over-suppression of translation and to restore homeostasis, the tRNA synthetase-mediated response also provides negative feedback mechanisms: The translation inhibition and stress-response genes activation effected by nuclear TyrRS would preserve ATP and amino acid and help alleviate the stressors that stimulate TyrRS nuclear translocation in the first place (Fig. 5). In addition, TyrRS also transcriptionally represses its own gene expression to ensure the repression is tapering off and not overdone. Interestingly, three other aaRS genes (i.e., WARS, SARS, and GARS) were also targeted by nuclear TyrRS (Fig. 2A), raising the possibility that TyrRS is not the only aaRS mediating this type of stress response. It is not hard to imagine that having multiple aaRSs involved in the regulation would increase the breadth (e.g., detecting the status of additional amino acids and tRNA species) and the capacity for the stress response. Interestingly, all 4 aaRSs targeted by TyrRS are stand-alone and not associated with the multi-synthetase complex (Bandyopadhyay and Deutscher, 1971; Hyeon et al., 2019), suggesting that mobility could be a required property for a synthetase to be involved in this integrated stress response network. In fact, nuclear functions have also been reported for TrpRS, SerRS, and GlyRS (Sajish et al., 2012; Shi et al., 2014; Yu et al., 2019). Particularly, TrpRS is highly induced by IFN-gamma, which also increases the amount of TrpRS in the nucleus (Sajish et al., 2012). Future studies should investigate nuclear functions of these and other aaRSs in stress response and whether the tRNA-controlled nuclear translocation is a common scheme.

Nuclear TyrRS-mediated translational suppression occurred at a later time point than eIF2a phosphorylation and mTOR inhibition during stress (Fig. 1A). In contrast to a sustained effect of H_2_O_2_ treatment on mTOR, as reflected by the decreased phosphorylation of 4E-BP1 and S6K, the effect of H_2_O_2_ treatment on eIF2a phosphorylation was acute due to a negative feedback regulation through ATF4 and its target gene GADD34, which dephosphorylates eIF2α to reinstate protein synthesis (Fig. 5). Indeed, we observed ATF4 activation in both normal and nuclear TyrRS deficient cells under oxidative stress (Fig. 1A). Interestingly, many protein synthesis genes suppressed by nuclear TyrRS, including YARS, WARS, SARS, and GARS, are also target genes of ATF4 with an opposite effect (activation) on their expression (Harding et al., 2003; Han et al., 2013) suggesting that nuclear TyrRS counteracts ATF4. Indeed, excluding TyrRS from the nucleus not only abolished the oxidative stress-induced repression of its target genes but also led to an over-stimulation of these genes (Fig. 2E), most likely mediated by the activated ATF4, no longer kept in check by nuclear TyrRS. Therefore, we envision that the entire cellular integrated stress response constitutes at least two stages, both aim to adapt and then to restore. The classical eIF2α/ATF4-based ISR provides the early response; if the stress is not yet resolved, the aaRS-based ISR mediates the late response, providing the cells with a second chance to adapt and survive through the stress condition. Moreover, the ATF4-mediated activation of aaRSs provides an internal connection and regulation between the two stages.

Aminoacyl-tRNA synthetase is the largest gene family associated with Charcot-Marie-Tooth disease (CMT), an inherited neurodegenerative disorder. The involvement of nuclear TyrRS in CMT has recently been demonstrated (Bervoets et al., 2019). CMT-causing mutations, including E196K, induce unique conformational changes in TyrRS (Blocquel et al., 2017), which confer the synthetase with the capacity to interact with transcriptional regulators such as TRIM28 and HDAC1 more strongly (Bervoets et al., 2019). Consistently, we show here that the E196K mutation enhances the inhibitory effect on target gene transcription and global protein synthesis. This observation is also consistent with an earlier report that CMT-mutations, including E196K in TyrRS, reduce global protein synthesis in *drosophila* CMT models in a manner independent of the enzymatic activity (Niehues et al., 2015). Importantly, this phenotype is not unique to TyrRS-CMT mutations but also can be induced by mutations in another CMT-linked tRNA synthetase (GlyRS), suggesting that a related nuclear function may apply to another aaRS. Although it is not yet clear whether impaired global translation is linked to the etiology of CMT, this observation does provide some *in vivo* evidence for a novel role of tRNA synthetase in translation regulation. Interestingly, *Drosophila-based* transcriptome analysis identified protein folding, regulation of cellular response to stress, and viral life cycle to be the top three gene ontology (GO) categories regulated by TyrRS (Bervoets et al., 2019). We speculate that dysregulation of the aaRS-based late integrated stress response network, including dysregulation of translation, could possibly contribute to the etiology of CMT.

Taken together, we describe here an alternative stress response pathway that operates distinctively from previously delineated regulations and allows cells to further adapt to stress after continuous exposure. The late integrated stress response ensures that translation is increased only upon resolution of stress and does not overshoot the levels of protein synthesis before the onset of stress. The ability to sense and initiate the response is inherent to the main component of this late integrated stress response, TyrRS, whose intrinsic properties as an aminoacyl-tRNA synthetase permit recognition of a broad range of inducers of cell stress.

## Material and Methods

### Cell lines and culture conditions

HEK-293 (human embryonic kidney) cells were purchased from ATCC. HEK-293 were grown in DMEM (Gibco) supplemented with 10% heat-inactivated fetal bovine serum (FBS, Gibco) and penicillin/streptomycin (Gibco). Cultures were maintained at 37°C in a humidified atmosphere under 5% CO_2_.

### Generation of ΔY/YARS and ΔY/YARS-NLS^Mut^ HEK293 cells

HEK293 cells were transfected with constructs pEco-Lenti-H1-shYARS-YARS or pEco-Lenti-H1-shYARS-YARS-NLS^Mut^ (Wei et al., 2014) which contain shRNA targeting 3’UTR of human YARS mRNA and coding sequence of human WT TyrRS or NLS-mutated (^242^NNKLNK^247^) TyrRS. About 48 hours after transfection, cells were selected under cell culture media supplemented with 1ug/ml puromycin for 10 days to generate ΔY/YARS and ΔY/YARS-NLS^Mut^ cells.

### Cell treatment

Cells were grown to 70-80% confluence before treatments. Transfections were performed using LipofectAMINE 2000 (Life Technology) according to the manufacturer’s instructions. For H_2_O_2_ (Millipore Sigma, H1009) treatment, H_2_O_2_ was diluted in PBS and added into the cell media to a final concentration of 200 μM for the indicated period of time. For homoharringtonine (HHT) treatment (Millipore Sigma, SML1091), HHT was added into cell media to a final concentration of 50 mM. For Trichostatin A (TSA, Millipore Sigma, T1952) treatment, TSA was added into cell culture media to a final concentration of 0.3 uM for 24 hours. For both HHT and TSA treatment, DMSO treatment was used in parallel as the control. For the E2F1knock down experiment, cells were transfected with shControl (Santa Cruz Biotechnology, sc-108060) or shE2F1 (Santa Cruz Biotechnology, sc-29297-SH) using LipofectAMINE 2000 for 72 hours.

### Surface sensing of translation (SUnSET)

Each sample size was one well of 6-wells plate, estimated around 300,000 cells per sample. Cells with the indicated treatment were incubated with 10 μg ml^-1^ of puromycin (Sigma Aldrich) for 30 min. After treatment, cells were washed twice in PBS, suspended in lysis buffer (Cell Signaling) and analyzed by Western blot with an antibody against puromycin (1:5000, Millipore, MABE343). a-Tubulin was used as a loading control. The amount of newly synthesized protein as detected by SUnSET was quantified using the ProteinSimple Western Blot analysis software and normalized to a-Tubulin signal. The ratio of both signals is shown.

### Western Blot assay

Each sample size was one well of 6-wells plate (except Co-IP experiments), estimated around 300,000 cells per sample. Cells were washed twice in phosphate-buffered saline (PBS), collected, suspended in lysis buffer (Cell Signaling) for 30 min, and centrifuged for 7 min at 12,000 * g; The soluble lysates were then collected, fractionated by SDS-PAGE, and transferred to PVDF membranes using the iBlot Dry Blotting System (Life Technologies). Membranes were blocked for 1 h with Tris Buffered Saline with Tween 20 (TBST) containing 5% nonfat dry milk. The following primary antibodies were diluted in 1% milk in TBST prior to usage at the indicated concentration: mouse monoclonal α-Tubulin (1:3000, Cell Signaling Technology, #3873), rabbit monoclonal p-p70 S6K (1:1000, Cell Signaling Technology, #9234), rabbit monoclonal p-4E-BP1 (1:1000, Cell Signaling Technology, #2855), rabbit monoclonal p70 S6K (1:1000, Cell Signaling Technology, #2708), rabbit monoclonal 4E-BP1 (1:1000, Cell Signaling Technology, #9644), rabbit polyclonal p-eIF2α (1:1000, Cell Signaling Technology, #9721), rabbit polyclonal eIF2α (1:1000, Cell Signaling Technology, #9722), rabbit monoclonal ATF-4 (1:1000, Cell Signaling Technology, #11815), rabbit polyclonal SARS (1:1000, made in-house), rabbit polyclonal WARS1 (1:3000, made in-house), mouse monoclonal GARS (1:4000, made in-house), rabbit monoclonal EEF1A1 (1:1000, Cell Signaling Technology, #3586), goat polyclonal RAE1 (1:1000, Abcam, ab36139), rabbit polyclonal HARS (1:1000, Abcam, ab137591), mouse monoclonal V5 (1:5000, Thermo Fisher Scientific, R960CUS), mouse monoclonal HDAC1 (1:1000, Cell Signaling Technology, #5356), mouse monoclonal HDAC2 (1:1000, Cell Signaling Technology, #5113), mouse monoclonal HDAC3 (1:1000, Cell Signaling Technology, #3949), mouse monoclonal RbAp46 (1:1000, NOVUS BIOLOGICALS, OTI5A4), mouse monoclonal RbAp48 (1:1000, NOVUS BIOLOGICALS, 13D10), mouse monoclonal CHD4 (1:1000, Abcam, ab70469), rabbit monoclonal MBD3 (1:1000, Abcam, ab157464), rabbit polyclonal MTA1 (1:1000, Abcam, ab71153), rabbit polyclonal p66a (1:1000, NOVUS BIOLOGICALS, NB100-56643), rabbit polyclonal LSD1 (1:1000, Abcam, ab17721), rabbit polyclonal TRIM28 (1:1000, Abcam, ab10483), rabbit polyclonal caspase 3 (1:1000, Cell Signalling Technology, #9662), rabbit polyclonal cleaved caspase 3 (1:1000, Cell Signaling Technology, #9661), rabbit monoclonal PARP (1:1000, Cell Signaling Technology, #9532), rabbit polyclonal SOD1 (1:1000, Abcam, ab13498), rabbit polyclonal Catalase (1:1000, Abcam, ab16731), rabbit polyclonal Thioredoxin (1:1000, Abcam, ab26320). After incubation with primary antibodies, the membranes were washed and incubated with HRP-conjugated anti-mouse or anti-rabbit secondary antibodies (1:3000, Cell Signaling Technology, #7076 or #7074) and followed by detection with ECL Western blot substrate (Thermo Fisher Scientific).

### Chromatin immunoprecipitation (ChIP) assay and sequencing

Each sample size was one 10 cm dish, estimated around 2,000,000 cells per sample. Cells were fixed with formaldehyde (1% final concentration) for 10 min at room temperature. The reaction was stopped by adding 125 mM glycine. ChIP assays were performed according to the protocol of ChIP-IT Express Enzymatic kit (Active Motif) using ChIP-grade antibody against V5 (Thermo Fisher Scientific, R960CUS), IgG (Cell signaling technology, #5415), H3K27Ac (Abcam, ab4729), HDAC1 (Abcam, ab7028), TRIM28 (Abcam, ab10483), or CHD4 (Abcam, ab70469). The eluted DNA fragments were subjected to library preparation and sequencing, or analyzed *via* a StepOnePlus Real-time PCR system using SYBR Select Master Mix (Applied Biosystems). For ChIP-qPCR, the primer sets targeting different genes were as follows: YARS ChIP (forward): 5’-TGA ATT ACA GTC ACT GTC AAA GCT ACA T-3’; YARS ChIP (reverse): 5’-AAT AGC CAA GTG CAG GGA GGT ACT GAG A-3’; SARS ChIP (forward): 5’-GGC AAC CGA ATC ATA TTC CTG TG-3’; SARS ChIP (reverse): 5’-CTC AAA CCG CTC TGC TTC CAA CT-3’; WARS ChIP (forward): 5’-GAG GGA GTG AAT TTC TTT GGT TGG-3’; WARS ChIP (reverse): 5’-GCA GCA GGG CTG GTT TAG GAT AG-3’; GARS ChIP (forward): 5’-CTG GCA TGA CAT TGT TTG GCT CTG TG-3’; GARS ChIP (reverse): 5’-CAG AGG ATA CCT GGC CTC CAC ATC AG-3’; HARS ChIP (forward): 5’-ACA TTA CAG GCA TGA GCC ACC-3’; HARS ChIP (reverse): 5’-CTA GCT GAT TGT GAT AAG AGA GAG AAC AGC-3’; EEF1A1 ChIP (forward): 5’-AGT TGC GTG AGC GGA AAG ATG-3’; EEF1A1 ChIP (reverse): 5’-CTA ATC GAG GTG CCT GGA CGG-3’; RAE1 ChIP (forward): 5’-GGG GCG GTT CAC GAT GTT-3’; RAE1 ChIP (reverse): 5’-CAG CAG TTT GCG AAT TGG TC-3’. The signal from IPed samples was quantified as percentage of the input from the whole cell lysate.

### TyrRS-Chip-seq

Each sample size was five 10 cm dish, estimated around 10,000,000 cells per sample. Libraries were prepared and sequenced by the Next Generation Sequencing Core at Scripps Research, La Jolla, California. Sequencing depth: V5 repeat 1: 20567024 (73.39% overall alignment to hg19), V5 repeat 1 input: 22802577 (92.61 % overall alignment), V5 repeat 2: 15358168 (80.43% overall alignment), V5 repeat 2 input: 23228662 (93.04% overall alignment), IgG control: 13431892 (78.67% overall alignment), IgG control input: 29488182 (89.09% overall alignment). Reads were quality filtered and adapter trimmed. In brief, the ENCODE 3 ChIP-Seq pipeline (ENCODE Project Consortium, 2012) was followed for read mapping and peak calling: reads were aligned to hg19 with Bowtie2 (Langmead and Salzberg, 2012) and duplicates were removed using picard tools and samtools (Li et al., 2009). TyrRS binding sites in the genome in comparison to a control input were identified using MACS (Zhang et al., 2008).

### Real-time PCR

Each sample size was one well of 6-wells plate, estimated around 300,000 cells per sample. Total RNAs were extracted from treated cells using Trizol (Life Technologies) according to the manufacturer’s instructions and then reverse-transcribed using the first strand cDNA synthesis kit (Thermo Fisher Scientific). The cDNA products were analyzed on a StepOnePlus Real-time PCR system using SYBR Select Master Mix (Applied Biosystems). The primers used for different target genes are: YARS RT (forward): 5’-CAA CCT GGC TGG ACT CGC GTG ACA GTT C-3’; YARS RT (reverse): 5’-AGG TTC CGG GTG ATA AGG TGC AGT TTC T-3’; SARS RT (forward): 5’-GGA ACC TTC TGC ACC CTT CTG TAC CCA T-3’; SARS RT (reverse): 5’-TTT CGC CTT CAA AGC CAT CTA CCA TCA C-3’; WARS RT (forward): 5’-CAA GGA CAT CAT CGC CTG TGG CTT TGA C-3’; WARS RT (reverse): 5’-CTG TGG GAA TGA GTT GCT GAA GGA GGG A-3’; GARS RT (forward): 5’-CTG TAG TTG CTC CAT TCA AAT GTT CCG TCC TC-3’; GARS RT (reverse): 5’-TCT CAT CAG TCC TGG CAT AGC GCC TTC C-3’; HARS RT (forward): 5’-GAA GAA TGA GAT GGT GGG AGA GA-3’; HARS RT (reverse): 5’-CAA TGC CAA ATA GGG TCA GGT A-3’; EEF1A1 RT (forward): 5’-ACA CGT AGA TTC GGG CAA GTC CAC CAC T-3’; EEF1A1 RT (reverse): 5’-TGA TAC CAC GTT CAC GCT CAG CTT TCA G-3’; RAE1 RT (forward): 5’-TGT GAT GAC TGG GAG CTG GGA TAA GAC T-3’; RAE1 RT (reverse): 5’-CAC CGA TGC TGA TGT TTC AGT GGA GAT T-3’.

### Nuclear-cytoplasmic fractionation

Each sample size was one 6cm dish, estimated around 500,000 cells per sample. Nuclear-cytoplasmic fractionation was conducted using the NE-PER Nuclear and Cytoplasmic Extraction Reagents kit (Thermo Fisher Scientific) according to the manufacturer’s protocol.

### TAP-mass spec

InterPlay Mammalian TAP System was purchased from Agilent (#240101). Human TyrRS (NM_003680.3) CDS was inserted into the Protein Interactions Expression Vector provided by the system to generate constructs to express TyrRS with SBP and CBP tags. The construct was transfected into HEK293 cells for 48 hours and then supernatant of whole cell lysate was added to the Streptavidin resin for SBP tag binding. The elution from the Streptavidin resin was next added to the Calmodulin resin for CBP tag binding. The product from the final elution and whole cell lysate supernatant were subjected to silver staining using Silver Stain Plus Kit (BIO-RAD, # 1610449) to check pull down efficiency and purity. The final eluted proteins were analyzed using the methods and protocols described in Pankow et al (Pankow et al., 2015) and Pankow et al (Pankow et al., 2016).

### Co-immunoprecipitation (Co-IP)

Each sample size was one 10 cm dish, estimated around 2,000,000 cells per sample. HEK293 cell lysates were prepared as described above for Western blot analysis. Protein G beads (Life Technologies) were pre-incubated with anti-TyrRS antibody for 30 min and then mixed with cell lysates overnight. The beads were then washed 3 times with buffer (100 mM NaCl, 50 mM Tris, pH 7.5, 0.1% Triton X-100, 5% glycerol), and immunoprecipitates were analyzed by SDS-PAGE and Western blot with the indicated antibodies.

### Cell viability assay

Each sample size was one well of 96-wells plate, estimated around 10,000 cells per sample. Relative cell viability was determined using a Cell counting kit-8 (Dojindo) and read on a microplate reader (Promega).

### Cellular ROS analysis

Each sample size was one well of 96-wells plate, estimated around 10,000 cells per sample. Cells were loaded with CM-H_2_DCFDA (Molecular Probe) for 30 min according to the manufacture’s instruction. Signals were read on a microplate reader (Promega).

### Statistical analysis

All data are presented as the mean ± s.e.m. Statistical significance between groups was evaluated using Student’s t-test. P < 0.05 was considered significant.

## Acknowledgements

The work was supported by US National Institutes of Health (R01 NS085092 and R01 NS113583 to X-.L. Yang, and P41 GM103533 to J.R. Yates) and the National Foundation for Cancer Research.

## Author Contributions

NW designed and performed experiments. HC, YS, GF, and NR conducted experiments and/or analyzed data. JRY and XLY designed and supervised experiments. NW, HC, and XLY wrote the manuscript.

## Corresponding author

Correspondence to Xiang-Lei Yang.

## Supplementary Figure Legends

**Supplementary Figure S1:**
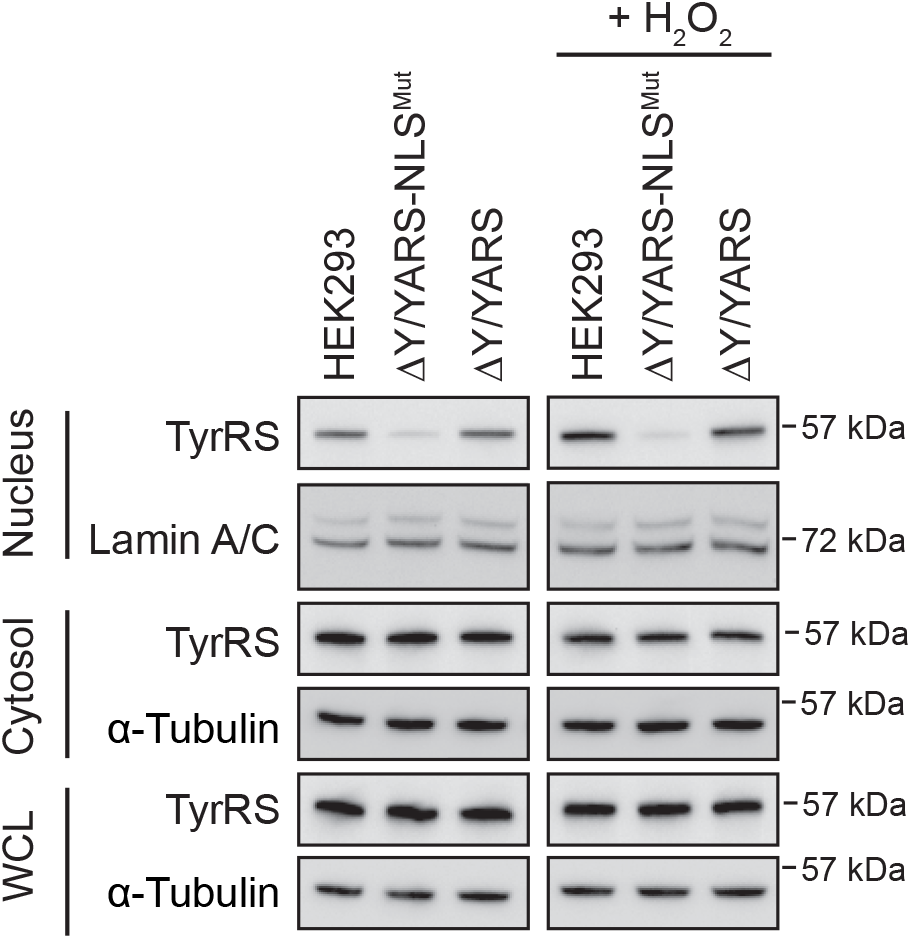
Cell fractionation and Western blot analysis to show that the expression level of TyrRS (WT or mutant) in ΔY/YARS and ΔY/YARS-NLS^Mut^ cells is similar to that of the endogenous TyrRS in the original, unmodified HEK293 cells and that ΔY/YARS-NLS^Mut^ cells are deficient in nuclear TyrRS with or without H_2_O_2_ treatment (12 hours). ΔY/YARS-NLS^Mut^: HEK293 cells with a knock down of endogenous TyrRS and expression of TyrRS with a mutated NLS (^242^KKKLKK^247^ to ^242^<u>NN</u>KL<u>N</u>K^247^). ΔY/YARS: HEK293 cells with a knock down of endogenous TyrRS and ectopic expression of wild type TyrRS. Lamin A/C: nuclear marker, α-Tubulin: cytoplasmic marker.

**Supplementary Figure S2:**
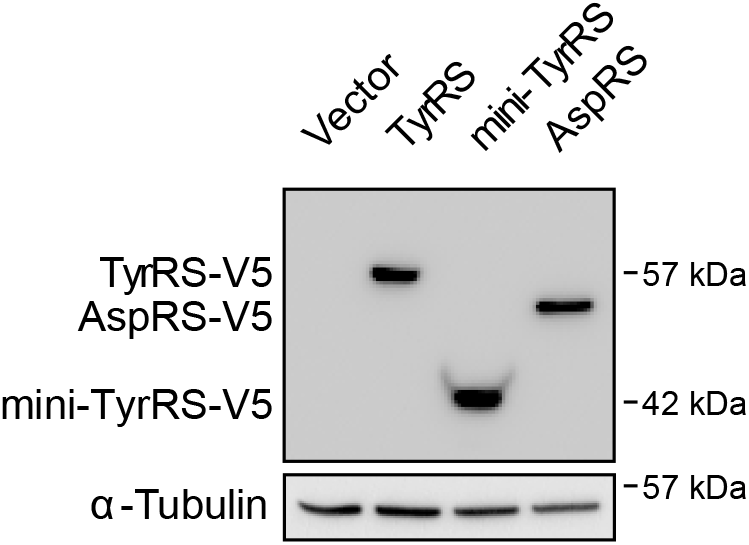
Western blot analysis to show the expression level of the V5-tagged TyrRS, mini-TyrRS and AspRS in HEK293 cells overexpressing each construct. Mini-TyrRS: Catalytic and tRNA binding domain of TyrRS.

**Supplementary Figure S3:**
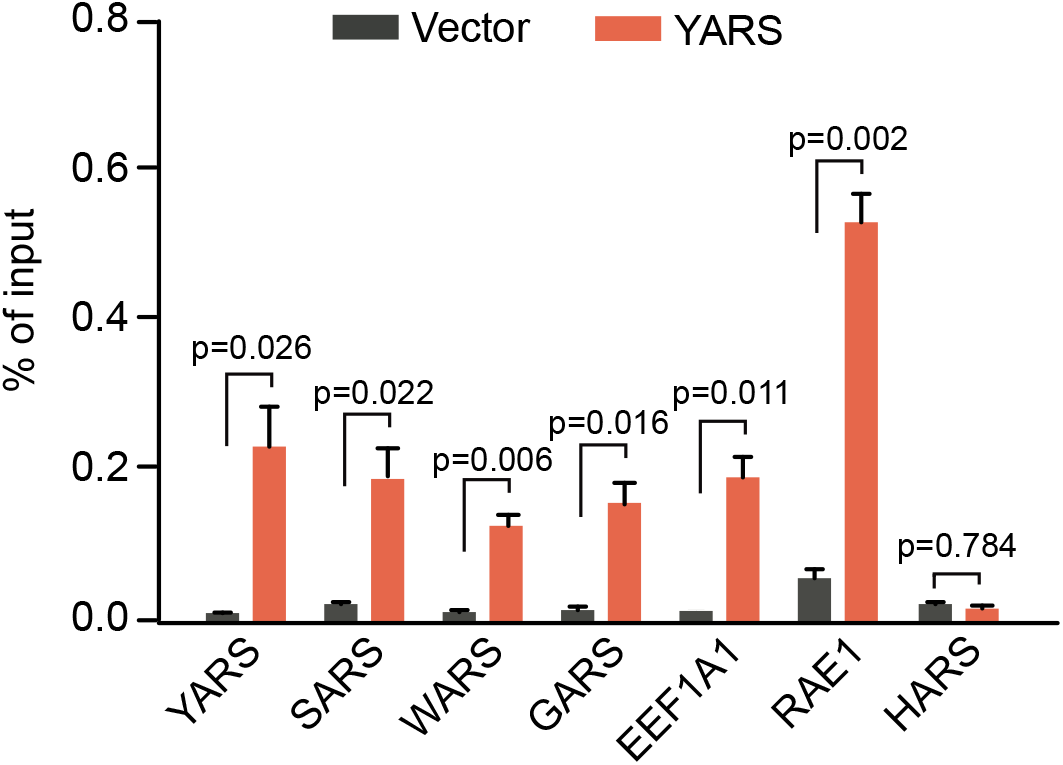
ChIP-qPCR assay confirms the DNA binding ability of TyrRS on translation-related genes. HEK293 cells transfected with TyrRS-V5 or Vector control were compared 24 hours after transgene overexpression by Chromatin IP using α-V5 antibody followed by RT-PCR analysis. n=3, biological replicates, one way Student’s t test. Vector: control, YARS: TyrRS overexpression.

**Supplementary Figure S4:**
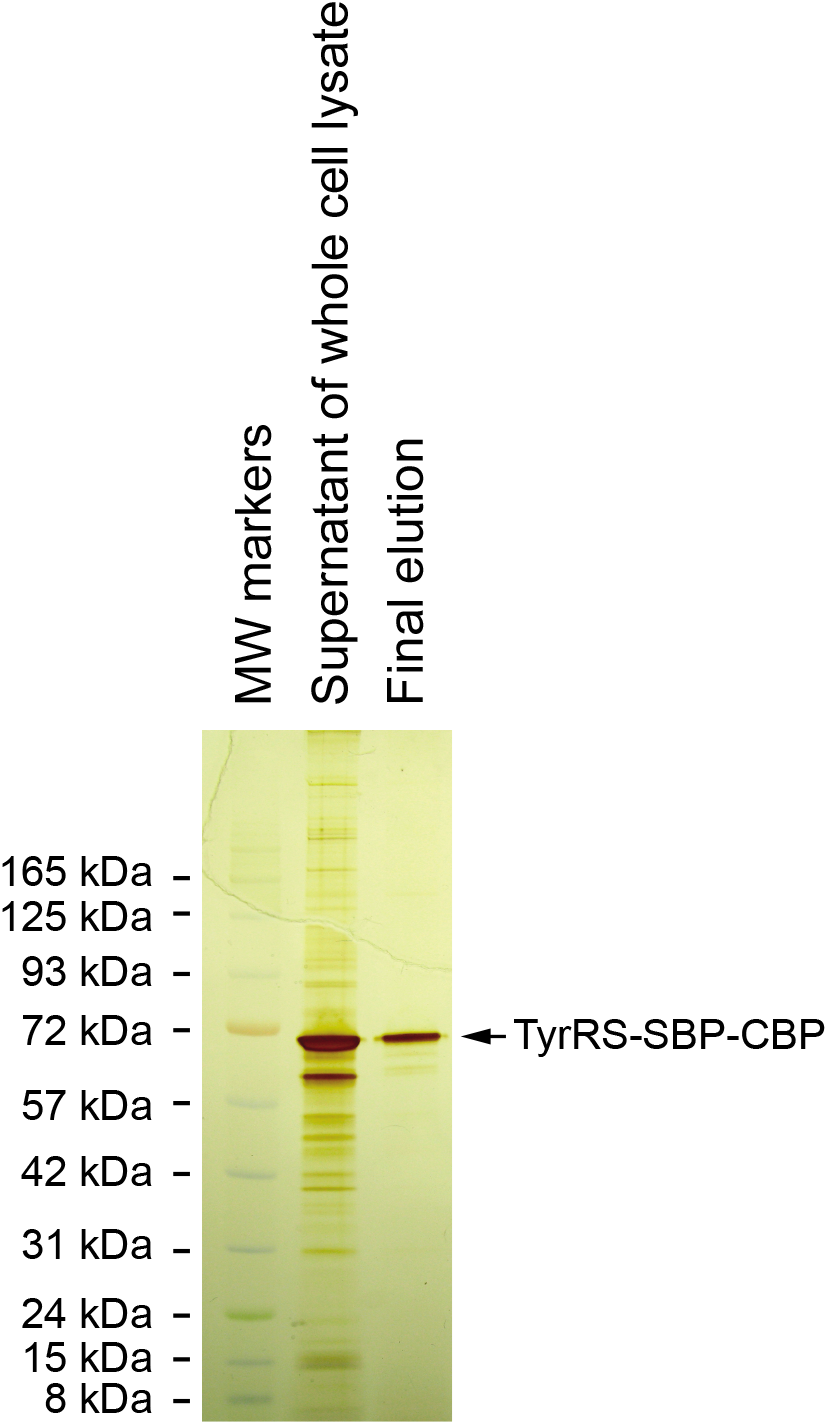
Silver staining showing the stringent Tandem Affinity Purification for isolating TyrRS and its interactome for mass spectrometry (TAP-MS) analysis. TyrRS-SBP-CBP: the fusion protein of TyrRS with a streptavidin binding peptide (SBP) and a calmodulin binding peptide (CBP).

**Supplementary Figure S5:**
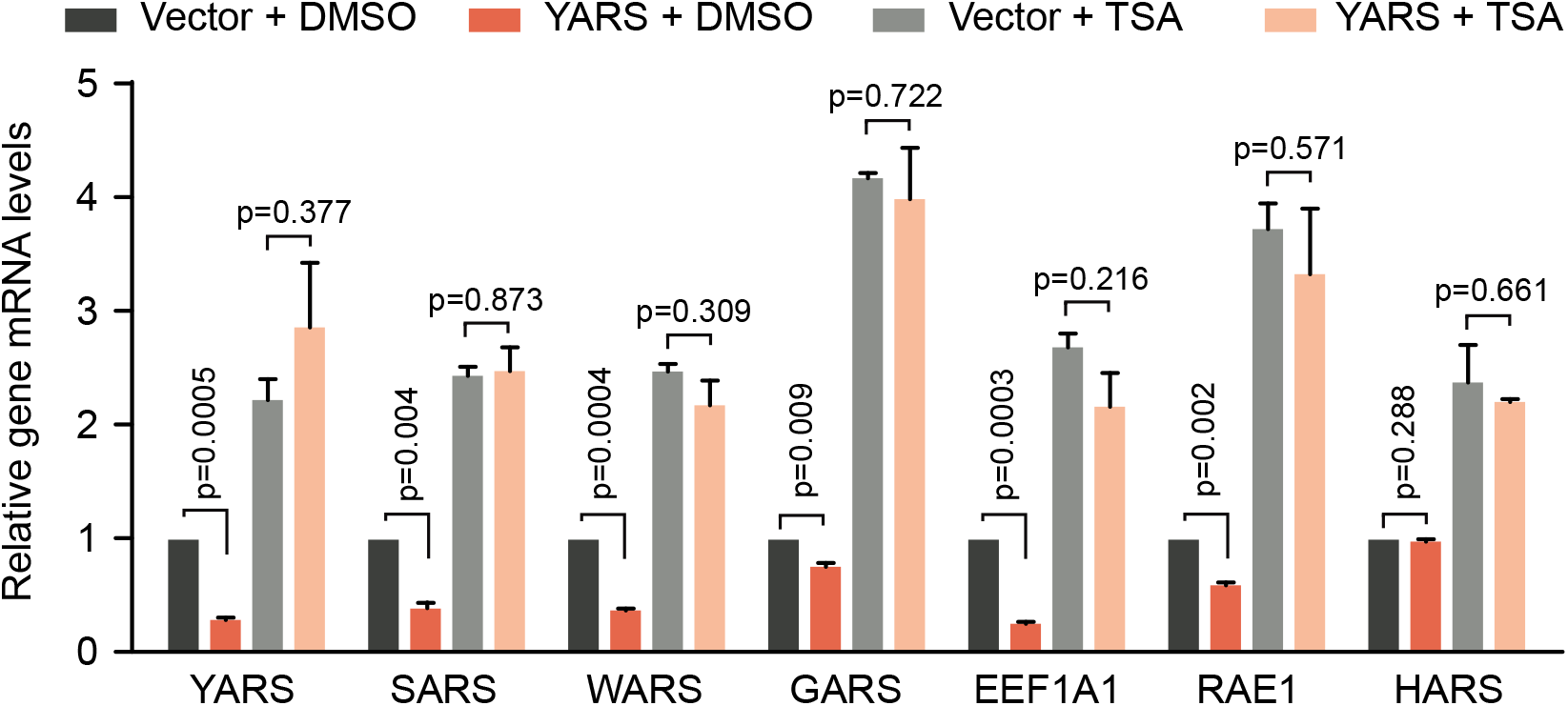
HDAC inhibitor TSA treatment blocks the inhibition effect of overexpressed TyrRS on its target gene transcription. Transcription of target genes were measured by RT-PCR using HEK293 cells with transgene overexpression for 24 hours, followed by DMSO or TSA treatment for another 24 hours. n=3, biological replicates, one way Student’s t test. Vector: control, YARS: TyrRS overexpression.

**Supplementary Figure S6:**
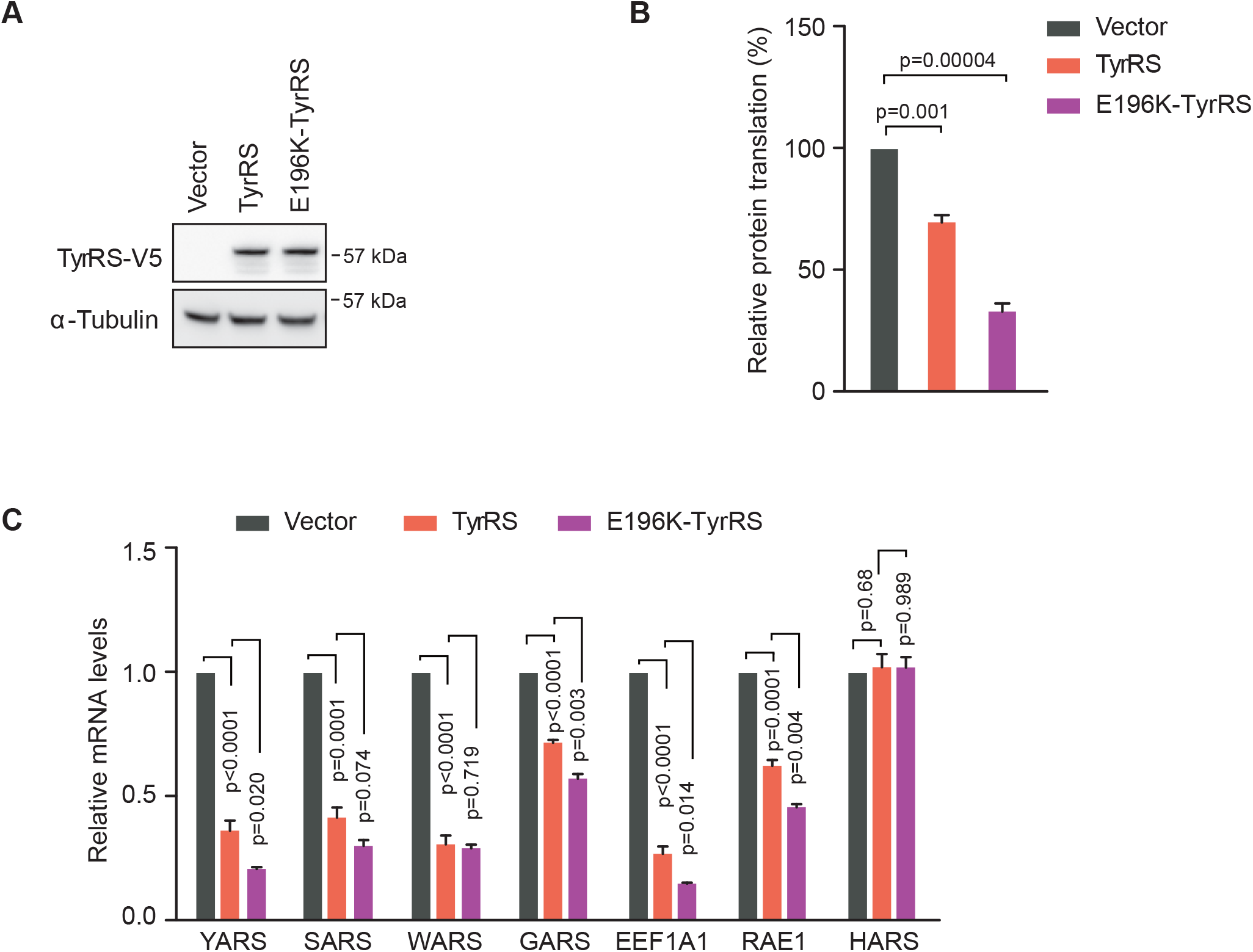
CMT-causing mutant E196K TyrRS shows a gain-of-function effect in suppressing protein synthesis and target gene expression compared to wild type TyrRS. A) Western blot analysis shows equal expression of WT and E196K TyrRS in HEK293 cells 24 hours after gene transfection. B) SUnSET analysis indicates a stronger effect of E196K TyrRS in protein synthesis inhibition compared to wild type TyrRS. n=3, biological replicates, one way Student’s t test. C) RT-PCR analysis indicates a stronger effect of E196K TyrRS in repressing the transcription of protein synthesis-related genes compared to the wild type TyrRS. n=3, biological replicates, one way Student’s t test.

**Supplementary Figure S7:**
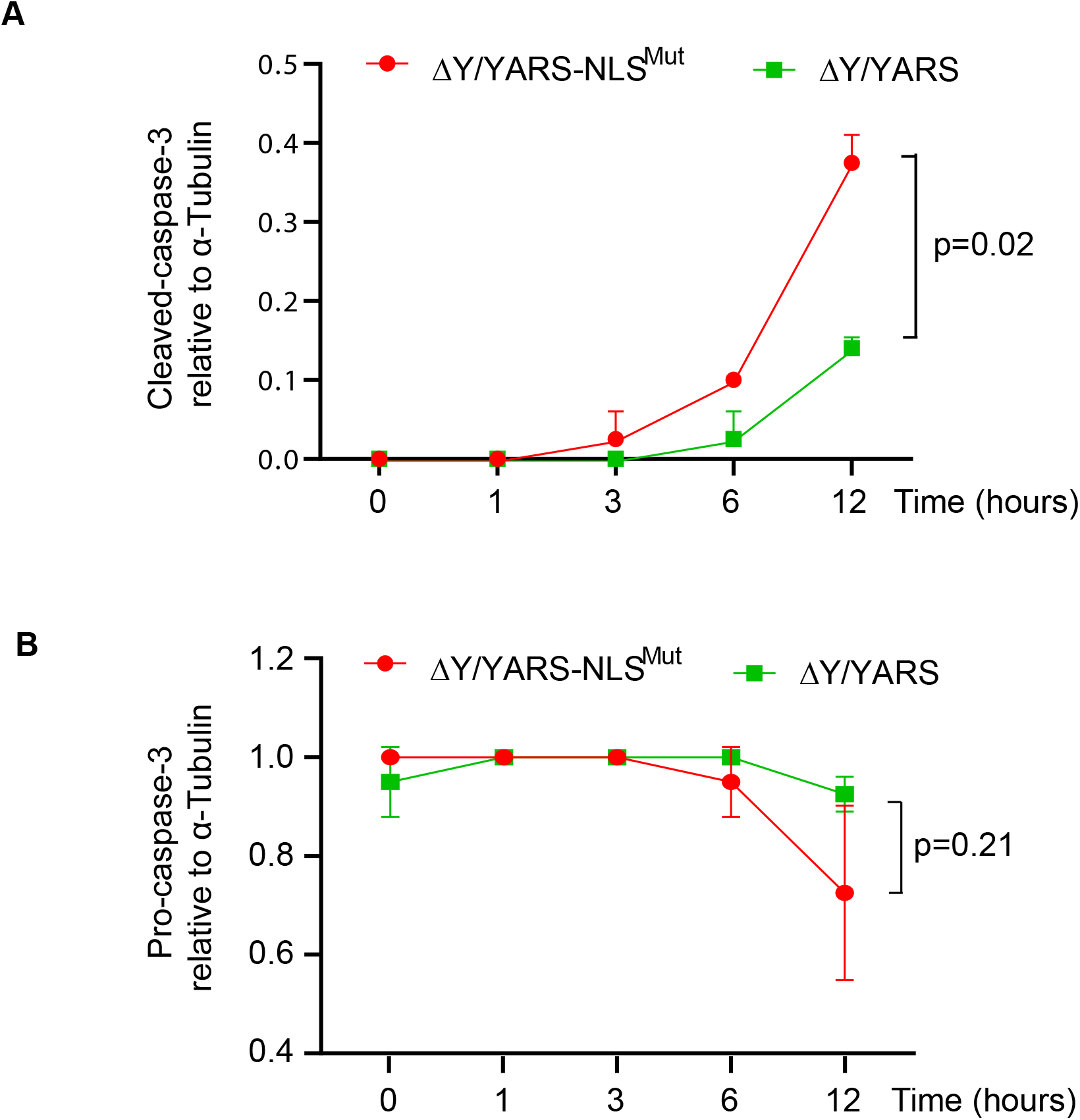
Quantification of caspase-3 cleavage as detected by Western blot analysis in Figure 4B. n=2, biological replicates, one way Student’s t test.

**Supplementary Figure S8:**
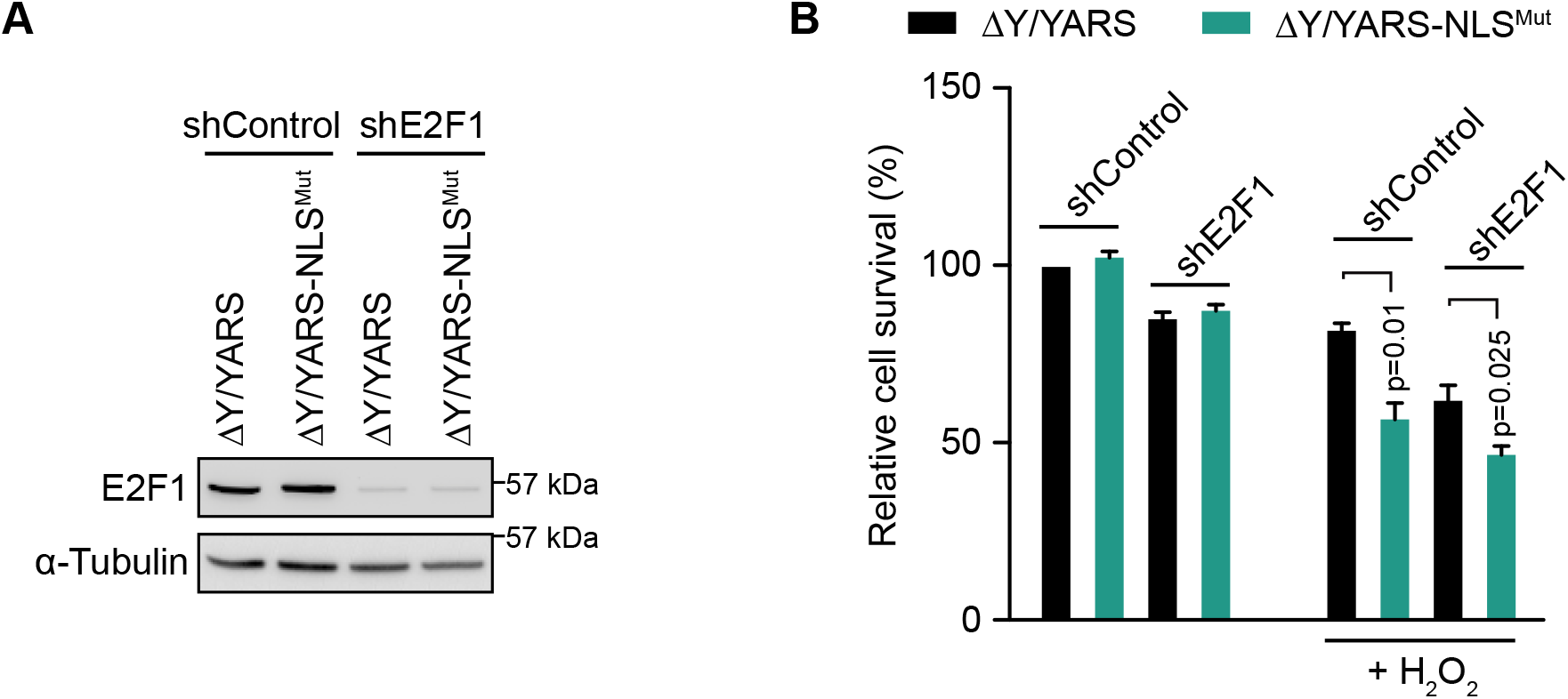
Nuclear TyrRS protects against cell death independent from its DNA damage protection effect, which is mediated by the transcription factor E2F1. ΔY/YARS-NLS^Mut^: HEK293 cells with a knock down of endogenous TyrRS and expression of TyrRS with a mutated NLS (^242^KKKLKK^247^ to ^242^<u>NN</u>KL<u>N</u>K^247^). ΔY/YARS: HEK293 cells with a knock down of endogenous TyrRS and ectopic expression of wild type TyrRS. A) Western Blot analysis confirms the knock down efficiency of shE2F1 in ΔY/YARS and ΔY/YARS-NLS^Mut^ cells. B) Stronger resistance to oxidative stress (H_2_O_2_ treatment for 36 hours) in cells with nuclear TyrRS (ΔY/YARS) compared to nuclear TyrRS-deficient cells (ΔY/YARS-NLS^Mut^) with or without E2F1 knockdown, indicating the protective effect of nuclear TyrRS against cell death is independent of E2F1. Relative cell viabilities were measured with a cell counting kit and viability of ΔY/YARS seeded at the same time but without H_2_O_2_ treatment was set at 100%. n=3, biological replicates, one way Student’s t test.

**Supplementary Figure S9:**
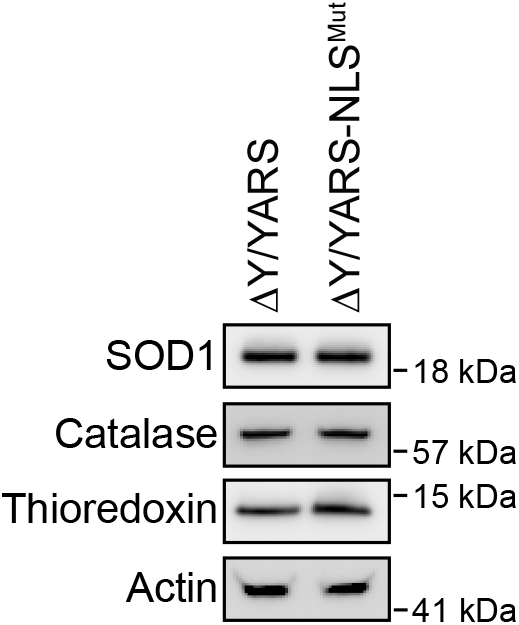
Nuclear TyrRS prevents cellular ROS over-accumulation independent of altering antioxidative stress response genes. Cells were treated with H_2_O_2_ for 36 hours followed by Western Blot analysis.

